# Analysis of reproductive traits associated with lipoprotein(a) deficiency in participants from the Pakistan Genome Resource

**DOI:** 10.64898/2026.11.05.764746

**Authors:** Christopher Koch, John E. Dominy, Allan M. Gurtan, Maleeha Zaman Khan, Shareef Khalid, Yoav E. Timsit, Juan P. Casas, Carolina Hache, Huilei Xu, Mike M. Mendelson, Ieuan Jones, Allison B. Goldfine, Danish Saleheen

## Abstract

**Aims:** Lipoprotein(a) [Lp(a)] is a genetically determined, independent risk factor for atherosclerotic cardiovascular disease, driving development of LPA-targeting therapies that markedly lower circulating Lp(a). Because *LPA*—the gene encoding apolipoprotein(a)—is absent in most model organisms, preclinical studies have limited value for assessing effects such as reproductive safety. Therefore, naturally occurring human *LPA* “knockouts” were analyzed to evaluate reproductive outcomes of on-target Lp(a) lowering.

**Methods and Results:** Individuals carrying high-confidence *LPA* loss-of-function (LoF) variants which reduce Lp(a) in a gene-dosage–dependent manner within the Pakistan Genome Resource were identified. LoF variants showed no depletion of homozygotes relative to Hardy–Weinberg expectations. *LPA* LoF associations with fertility-related traits (ever having a child and number of children) under additive and recessive models were assessed using regenie-based burden testing, including sex-stratified analyses, and no significant associations were observed. A recall-by-genotype study of female LoF homozygotes alongside heterozygous and non-carrier family member controls was performed, surveying reproductive histories and offspring outcomes. Across 107 pregnancies and 87 children in 19 homozygous LoF women, rates of miscarriage, maternal complications (including preeclampsia, hypertension, and gestational diabetes), delivery complications, neonatal intensive care admission, birth defects, and child health problems were comparable across genotypes with no statistically significant genotype–outcome associations.

**Conclusions:** These genetic and by-genotype analyses indicate that complete lifelong *LPA* deficiency due to high-confidence LPA loss-of-function variants is not associated with overt adverse maternal fertility, pregnancy or child health outcomes. These findings offer human genetic reassurance regarding reproductive and developmental safety in the context of sustained Lp(a)-lowering, with implications for long-term therapies targeting Lp(a).

**Translational Perspective:** Despite advances in clinical management of atherosclerosis, cardiovascular event rates remain high and novel therapeutic approaches to reduce residual risk are needed. Lp(a) is an independent, genetic risk factor for atherosclerosis. Lp(a)-lowering therapies are under development. Prior to marketing reproductive safety is typically evaluated. However, rodents do not express apolipoprotein(a), the unique Lp(a) structural protein, rendering them unsuitable for evaluating on-target reproductive safety. This human genetics study provides evidence that lifelong, complete LPA deficiency caused by naturally occurring loss-of-function variants is not associated with reduced fertility, adverse pregnancy outcomes, congenital abnormalities, or impaired child health. These findings provide human genetic support for reproductive and developmental safety of chronic Lp(a)-lowering therapies and the approach may inform evaluations of other therapeutic targets.

## Introduction

Lipoprotein(a) [Lp(a)] is a lipid particle that contains a unique glycoprotein, apolipoprotein(a) (apo(a)), covalently linked to an LDL-derived particle containing apolipoprotein B (apoB). Lp(a) is a genetic, independent, and likely causal risk factor for atherosclerotic cardiovascular disease (ASCVD), based on epidemiological and Mendelian randomization (MR) studies demonstrating that elevated Lp(a) increases ASCVD risk^1–4^. Proposed mechanisms include increased oxidative phospholipid content, increased deposition of cholesterol in arterial walls, a proinflammatory vascular environment, perturbations in thrombostasis, and calcific aortic valve stenosis^5^.

Lp(a) concentrations are largely genetically determined and remain stable over life^6^. Diet, exercise, and weight loss do not meaningfully lower Lp(a) in most individuals. Three to 5% of the population have null *LPA* variants, and even lifelong absence of Lp(a) appears to have no major adverse phenotypes, although some but not all studies suggest an association of low Lp(a) with type 2 diabetes^7^. MR studies supporting causality have driven development of therapeutics, including antisense oligonucleotides (ASOs) and small interfering RNAs (siRNAs), and small molecules to durably lower Lp(a) concentrations, which lower apo(a) and Lp(a) by up to 80% (ASOs)^8^ and over 90% (siRNAs)^9^.

A challenge to developing novel therapies that lower Lp(a) is that the common preclinical species lack orthologs of human *LPA*; expression is limited to humans, Old World monkeys, apes, and hedgehogs. Sequence divergence between human and non-human primates (NHPs) further limits translatability.

Human genetics can de-risk potential long-term, on-target toxicities^10^. Genetic variants that mimic pharmacological intervention – particularly loss-of-function (LoF) variants – can approximate chronic drug effects. Identification of homozygous LoF carriers informs consequences of complete protein deficiency^11^. Reproductive outcomes are of particular interest to inform use in women of childbearing potential given the broad population who may receive Lp(a) lowering therapies.

Here, we report the identification and characterization of the reproductive health outcomes of homozygous *LPA* LoF women from the Pakistan Genome Resource (PGR), a South Asian biobank of approximately 173K whole exome/whole genome sequenced individuals enriched for consanguinity rare LoF homozygotes^12^. *LPA* LoF homozygotes and consenting family members were recontacted, genotyped and retrospectively assessed for pregnancy, maternal, fetal and postnatal outcomes. Homozygous *LPA* LoF was not associated with differences in maternal or child health for assessed traits versus controls.

## Methods

### Pakistan Genome Resource (PGR) overview

The PGR consists of 166,625 whole exomes and 6,678 whole genomes from 173,303 Pakistani participants (38.8% female) recruited from public and private hospitals in 23 cities across Pakistan, covering most major ethnicities present in Pakistan. The PGR encompasses nested case-control recruitments conducted for a variety of diseases, including myocardial infarction, diabetes, heart failure, metabolic dysfunction associated steatohepatitis, and chronic kidney disease^13,14^. For each participant, information was collected on demographic factors, lifestyle, medical and family history, and anthropometry. Biological samples were collected and stored, including blood derived-DNA, plasma, and serum for sequencing and biomarker analysis.

### Sequencing of the Pakistan Genome Resource (PGR) cohort

Exome sequencing of the PGR biobank was performed in multiple batches at two different locations: the Broad Institute^14^ and the Regeneron Genetics Center (RGC)^15,16^. Whole genome sequencing was performed within the Trans-Omics for Precision Medicine (TOPMed) consortium^17^.

#### PGR sample quality control (QC), variant QC, and variant annotation

Samples were excluded if there was discordance between their reported and computationally predicted sex, or if they were predicted to be genetic duplicates (including duplicate samples from the same individual or possibly identical twins) within or across sequencing batches. In the RGC datasets, samples were additionally excluded if fewer than 80% of bases were covered at 20x or >5% human contamination was observed^18^. For a subset of the RGC datasets, whole plates were excluded if self-reported gender was discordant with computationally predicted sex for > 10% of samples. After filtering, a total of 166,625 whole exome sequences (WES) and 6,678 whole genome sequence (WGS) samples were analyzed. Variant-level QC was applied separately to each sequencing batch. Before applying QC filters, multiallelic sites were decomposed and all variants were left-aligned and normalized. In the WES dataset sequenced at the Broad Institute, individual variant calls were set to missing if they did not meet all of the following criteria: Quality by Depth (QD) ≥ 2 for indels, or QD ≥ 3 for single nucleotide variants (SNVs); Variant Quality Score by Recalibration (VQSR) = “PASS” or (VQSR = “VQSRTrancheSNP99.60to99.80” and allele-count = 1); 0.25 ≤ allele-balance ≤ 0.75 for heterozygous calls, or allele-balance ≥ 0.9 for homozygous calls; depth (DP) ≥ 10. For a batch of WGS data sequenced at the Broad Institute, calls with DP <10 were excluded. In the datasets sequenced at the RGC, individual variant calls were set to missing if they did not meet all of the following criteria: 0.25 ≤ allele-balance ≤ 0.75 for heterozygous calls, or allele-balance ≥ 0.9 for homozygous calls; depth (DP) ≥ 7 for SNVs, or DP ≥ 10 for indels. After applying the above variant QC steps, variants were excluded entirely if they had a call rate < 0.9. All variant QC steps were performed using Hail. Variants were annotated using Variant Effect Predictor (VEP) version 107^19^ and Loss-of-Function Transcript Effect Estimator (LOFTEE)^19,20^. All variant annotations are reported relative to the Ensembl canonical transcript, version 107.

### Pakistan Genome Resource (PGR) analysis

Burden analyses in PGR were used to assess the relationship between *LPA* LoF and Lp(a) levels, number of children, and whether an individual has had any children. All burden analyses were performed using regenie^21^: v3.1.3 for Broad Institute batches and v3.1.1 on the DNAnexus Apollo platform for RGC batches. Regenie results from each sequencing platform were meta-analyzed using inverse variance weighted meta-analysis as implemented in METAL (2011-03-25 release)^22^. Quantitative traits were transformed by the rank-based inverse standard normal function, applied within each sequencing batch by regenie. Effect sizes are reported in standard deviations of the normalized trait. Serum Lp(a) concentrations were measured using an immunoturbidimetric assay (Denka Seiken, San Jose, CA, USA) ^23^. Lp(a) concentrations and number of children were independently analyzed for association with *LPA* genotype using both additive and recessive linear regression models as implemented in regenie. Having had any children was analyzed with both additive and recessive logistic regression models with Firth fallback as implemented in regenie. All analyses with the PGR biobank data, which includes both men and women, were adjusted for age, sex, age*sex, age^2^ and the first 10 genetic principal components. Genetic principal components were calculated within each sequencing batch. Prior to principal component analysis (PCA), variants were first filtered to minor allele frequency > 1% and then LD-pruned using the PLINK (v2.00a3.7) flag ‘--indep-pairwis’ with a window size of 200kb and an r^2^ threshold of 0.5^24^. PCA was performed on this pruned dataset using PC-AiR, as implemented in the GENESIS package (version 2.36.0)^25^.

### Recall by genotype study

Using whole exome and whole genome sequence data from the PGR biobank, 157 female homozygous carriers of the 3 most common *LPA* LoF variants (rs41272114, rs559580002, and rs533200556; see **Supplemental Table 1**) and 157 female controls matched for age, autozygosity, self-reported ethnicity, and case/control status, and not carrying any *LPA* variant with minor allele frequency < 5% were selected for recontact through the Center of Non-Communicable Diseases in Karachi, Pakistan. Specifically, for each of the 157 female homozygous carriers, individuals of the same self-reported ethnicity, and, wherever possible, same case/control recruitment status were selected as candidate controls. Among candidate controls, dissimilarity to the proband was calculated as the sum of the squared differences in z-scores for autozygosity and age. The individual with minimum z-scored-based dissimilarity was selected. The study was conducted under the Institutional Review Board (IRB) approval of IRB Committee of Center for Non-Communicable Diseases (NIH registered IRB 00007048, IORG0005843 and FWAS00014490).

After obtaining consent from the proband and family members, questionnaires regarding past medical and family history were administered by trained research staff in the local language. Questions included collection of both quantitative and non-quantitative data. Responses related to open ended medical history were tabulated. Recruitment proceeded until all potentially recontact-able individuals either consented or declined to participate. Sex was self-reported. Questions related to reproductive health are available in **Supplement**. Physical anthropometric information, and blood and urine samples were collected with details available in **Supplement**.

#### Statistical analysis

The number of pregnancies and the number of children per woman were analyzed using linear regression with age as a covariate. Pregnancy outcomes (livebirth, stillbirth, miscarriage, maternal and delivery complications) were assessed using mixed effects logistic regression with participant ID as a random intercept and *LPA* LoF genotype as a fixed effect.

Child outcomes (sex, neonatal intensive care unit admission, having birth defects, survival to 6 months, 1 year, and 5 years) were analyzed using mixed effects logistic regression with maternal ID as a random intercept and maternal *LPA* LoF genotype as a fixed effect. Additive and recessive models were applied for all outcomes.

Linear and logistic regression were performed using statsmodels (Python module version 0.14.5) and mixed effects logistic regression using the *glmer* function of the *lme4* R package, via *rpy2* from Python. Nominal p-values unadjusted for multiple testing are reported.

Models for stillbirth showed “boundary (singular) fit” warnings and were not reported. The recessive model for abortion failed to converge due to no homozygous LoF cases and was not reported.

#### Isolation of genomic DNA

DNA was extracted from leukocytes of peripheral whole blood using a salting out method. DNA concentrations were determined by ultraviolet quantification a using Nanodrop™ 2000 Spectrophotometer (Thermo Scientific™, USA).

#### Sanger sequencing

Whole blood-derived DNA from recalled participants was used to assess carrier status and zygosity for variants of interest via Sanger sequencing. Primer design, PCR reaction conditions, and sequencing are described below.

#### Primer design

Primers were designed for the 3 variants targeted for recall by genotype studies using Primer 3Plus Software^26^, and *in-silico* PCR was run using the UCSC Genome Browser’s tool (see **Supplemental Table 1** for primer sequences).

#### Polymerase chain reaction (PCR) and product cleanup

The target DNA region was amplified using AccuStart™ II PCR SuperMix (QuantaBio, USA). The product was run on 2% agarose gel to confirm a single intact band, then purified using ExoSAP-IT Express PCR Product Cleanup reagent (Thermo Scientific, USA), per manufacturer’s instructions.

#### Sanger sequencing and analysis

The PCR product was used for Sanger sequencing. Cycle sequencing was performed using BigDye™ Terminator v3.1 Cycle Sequencing Kit, and extension products were purified using the BigDye XTerminator™ Purification Kit (all by Thermo Scientific, USA). The purified sequencing reactions were analyzed on an SeqStudio™ Genetic Analyzer (Applied Biosystems, Thermo Scientific, USA).

## Results

### Identification of *LPA* loss-of-function (LoF) homozygotes in the Pakistan Genome Resource

PGR includes participants with high-confidence predicted LoF variants in the *LPA* gene, including those disrupting splice sites, causing frameshifts, or introducing premature stop codons (**Table 1**). Thirteen of these variants have at least 1 homozygous carrier. Among 26 *LPA* LoF homozygotes, the highest measured Lp(a) was 6.1 mg/dl, observed in a carrier of two alleles of the splice donor variant, c.4289+1G>A (rs41272114). A burden analysis of Lp(a) concentrations broken down by zygosity shows high confidence *LPA* LoF variants produce a gene-dosage reduction in circulating Lp(a), confirming they are LoF at the protein level (β=-0.74, 95% CI [-0.80, -0.67], P=2.1e-211; **Figure 1**).

**Figure 1:**
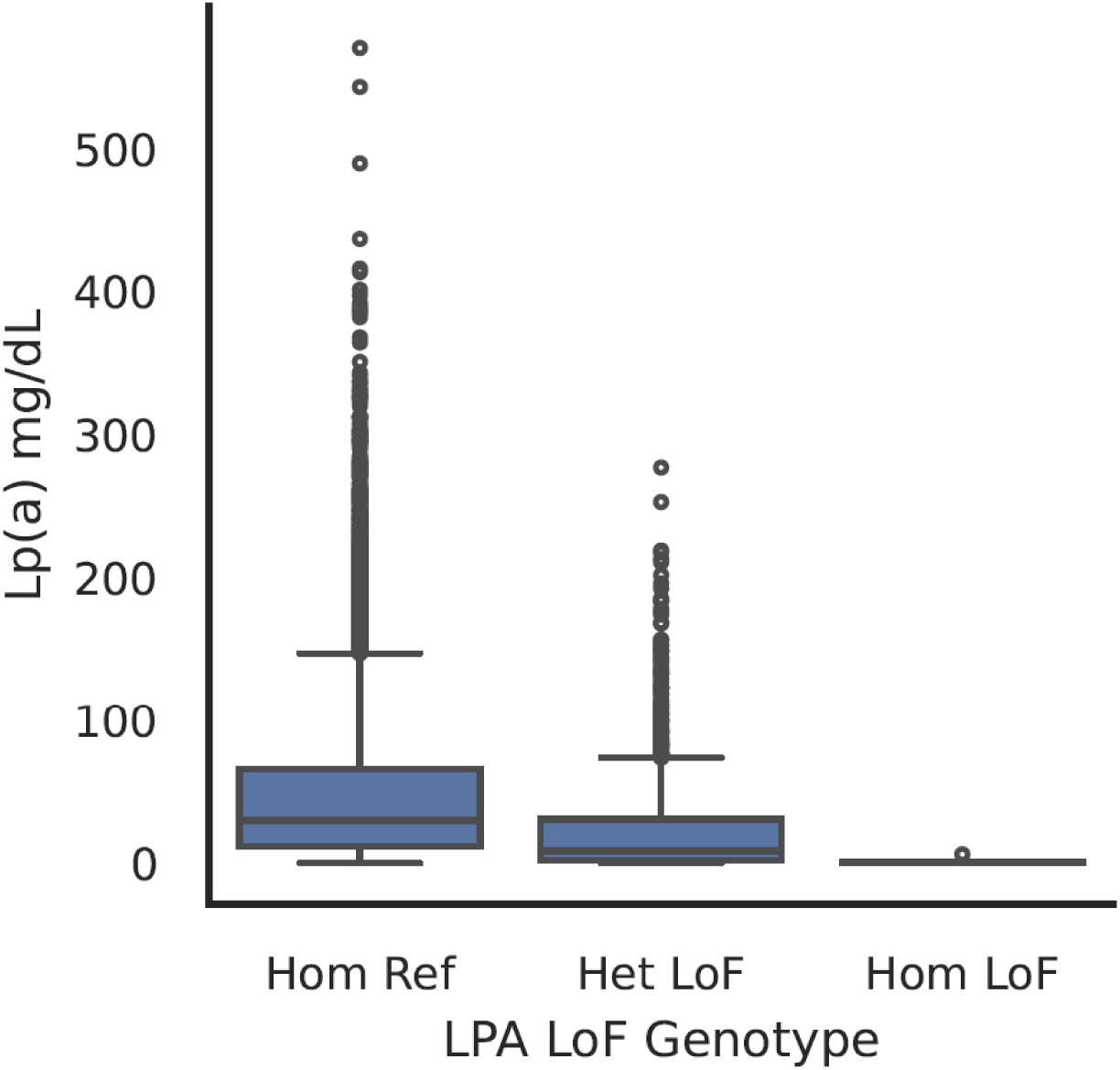
High confidence LPA loss of function (LoF) variants produce a gene-dosage reduction in circulating Lp(a).

**Table 1.** PGR *LPA* high-confidence LoF variant statistics. Descriptive annotations, allele counts and frequencies, Hardy-Weinberg equilibrium (HWE) statistics, and sample summary statistics by variant genotype for all high-confidence LoF *LPA* variants in PGR. CPRA, chromosome:position:reference_allele:alternative_allele; HGVSc, Human Genome Variation Society coding change; HGVSP Human Genome Variant Society protein change; AF, minor allele frequency; HomRR, homozygous reference individual count; HetRA, heterozygous individual count; HomAA, homozygous LoF individual count.

| CPRA | rsID | HGVSc | HGVSp | AF | HomRR | HetRA | HomAA | Expected HomAA Under HWE | HWE P-value | % Female<br>(HomRR\|HetRA\|HomAA) | Mean Age<br>(HomRR\|HetRA\|HomAA) | Mean Female Age<br>(HomRR\|HetRA\|HomAA) | Mean Male Age<br>(HomRR\|HetRA\|HomAA) | Mean Number of Children | Mean Number of Children to Females | Mean Number of Children to Males |
|---|---|---|---|---|---|---|---|---|---|---|---|---|---|---|---|---|
| chr6:160585045:C:T | rs41272114 | c.4289+1G>A | n/a | 0.0367 | 160865 | 11896 | 404 | 233.0027 | 0.00E+00 | 38.80\|38.69\|38.37 | 50.28\|50.44\|50.07 | 49.04\|49.35\|48.25 | 51.07\|51.12\|51.21 | 4.43\|4.42\|4.36 | 4.70\|4.71\|4.39 | 4.25\|4.22\|4.34 |
| chr6:160611561:C:T | rs559580002 | c.2603+1G>A | n/a | 0.0062 | 170944 | 2057 | 45 | 6.6595 | 0.00E+00 | 38.78\|38.94\|31.11 | 50.30\|49.95\|49.40 | 49.06\|48.94\|51.50 | 51.08\|50.59\|48.45 | 4.42\|4.53\|4.72 | 4.70\|4.71\|5.45 | 4.25\|4.41\|4.43 |
| chr6:160650337:C:A | rs533200556 | c.209+1G>T | n/a | 0.0016 | 172591 | 528 | 10 | 0.4336 | 0.00E+00 | 38.79\|36.93\|40.00 | 50.30\|49.49\|46.60 | 49.07\|48.72\|54.25 | 51.08\|49.95\|41.50 | 4.42\|4.24\|3.88 | 4.70\|4.39\|5.00 | 4.25\|4.15\|3.50 |
| chr6:160542737:A:AC | rs752960449 | c.5469dup | p.Cys1824ValfsTer72 | 0.0009 | 172883 | 305 | 3 | 0.1396 | 1.83E-14 | 38.79\|40.00\|33.33 | 50.29\|50.23\|50.33 | 49.06\|49.17\|36.00 | 51.07\|50.94\|57.50 | 4.42\|4.26\|4.33 | 4.70\|4.56\|6.00 | 4.25\|4.06\|3.50 |
| chr6:160585206:C:G | rs199536939 | c.4130-1G>C | n/a | 0.0006 | 173096 | 189 | 1 | 0.0526 | 3.60E-05 | 38.79\|44.44\|0.00 | 50.29\|49.41\|38.00 | 49.06\|47.38\|n/a | 51.07\|51.04\|38.00 | 4.42\|4.68\|4.00 | 4.70\|5.03\|n/a | 4.25\|4.40\|4.00 |
| chr6:160577135:C:T | rs200099994 | c.4631+1G>A | n/a | 0.0003 | 172600 | 99 | 1 | 0.0148 | 5.55E-16 | 38.77\|36.36\|0.00 | 50.30\|52.84\|57.00 | 49.07\|51.44\|n/a | 51.07\|53.63\|57.00 | 4.42\|4.45\|6.00 | 4.70\|4.74\|n/a | 4.25\|4.28\|6.00 |
| chr6:160548544:CTG:C | rs754796697 | c.5087_5088del | p.Thr1696ArgfsTer21 | 0.0002 | 35677 | 13 | 0 | 0.0012 | 9.73E-01 | 19.48\|46.15\|n/a | 53.67\|50.62\|n/a | 54.71\|50.50\|n/a | 53.42\|50.71\|n/a | 4.35\|5.17\|n/a | 4.78\|5.20\|n/a | 4.25\|5.14\|n/a |
| chr6:160577168:C:T | rs773026456 | c.4599G>A | p.Trp1533Ter | 0.0002 | 173238 | 52 | 0 | 0.0039 | 9.50E-01 | 38.80\|26.92\|n/a | 50.29\|47.85\|n/a | 49.06\|44.07\|n/a | 51.07\|49.24\|n/a | 4.42\|3.74\|n/a | 4.70\|4.08\|n/a | 4.25\|3.62\|n/a |
| chr6:160606577:C:T | rs760239476 | c.2685G>A | p.Trp895Ter | 0.0002 | 173210 | 81 | 1 | 0.0099 | 0.00E+00 | 38.80\|38.27\|0.00 | 50.29\|50.40\|45.00 | 49.06\|48.87\|n/a | 51.07\|51.34\|45.00 | 4.42\|4.55\|7.00 | 4.70\|4.50\|n/a | 4.25\|4.57\|7.00 |
| chr6:160540069:TGC:T | rs1382955417 | c.5707_5708del | p.Ala1903ArgfsTer15 | 0.0001 | 137530 | 40 | 0 | 0.0029 | 9.57E-01 | 43.81\|52.50\|n/a | 49.41\|49.08\|n/a | 48.41\|47.81\|n/a | 50.20\|50.47\|n/a | 4.45\|4.14\|n/a | 4.69\|4.33\|n/a | 4.25\|3.94\|n/a |
| chr6:160540073:T:TTC | rs1454111968 | c.5704_5705insGA | p.Gln1902ArgfsTer8 | 0.0001 | 99989 | 23 | 0 | 0.0013 | 9.71E-01 | 47.04\|52.17\|n/a | 47.58\|48.04\|n/a | 46.92\|44.50\|n/a | 48.17\|51.91\|n/a | 4.43\|4.55\|n/a | 4.65\|5.10\|n/a | 4.22\|4.00\|n/a |
| chr6:160548661:T:C | rs143431368 | c.4974-2A>G | n/a | 0.0001 | 173134 | 43 | 1 | 0.0029 | 0.00E+00 | 38.79\|32.56\|0.00 | 50.29\|51.14\|23.00 | 49.06\|51.29\|n/a | 51.07\|51.07\|23.00 | 4.42\|4.26\|n/a | 4.70\|4.50\|n/a | 4.25\|4.15\|n/a |
| chr6:160556125:G:A | rs749929484 | c.4873C>T | p.Arg1625Ter | 0.0001 | 173251 | 41 | 1 | 0.0027 | 0.00E+00 | 38.80\|39.02\|0.00 | 50.29\|48.20\|46.00 | 49.06\|51.00\|n/a | 51.07\|46.40\|46.00 | 4.42\|4.24\|5.00 | 4.70\|4.80\|n/a | 4.25\|3.86\|5.00 |
| chr6:160578522:C:A | rs751547669 | c.4471+1G>T | n/a | 0.0001 | 144226 | 13 | 2 | 0.0005 | 0.00E+00 | 43.08\|61.54\|50.00 | 49.58\|48.62\|65.50 | 48.52\|48.00\|56.00 | 50.38\|49.60\|75.00 | 4.42\|4.27\|6.00 | 4.67\|4.57\|1.00 | 4.23\|3.75\|11.00 |
| chr6:160595537:T:C | rs776068781 | c.3288-2A>G | n/a | 0.0001 | 6675 | 1 | 0 | 0 | 9.95E-01 | 27.99\|100.00\|n/a | 53.02\|62.00\|n/a | 52.17\|62.00\|n/a | 53.35\|n/a\|n/a | 3.99\|5.00\|n/a | 4.27\|5.00\|n/a | 3.88\|n/a\|n/a |
| chr6:160605045:C:T | rs201480327 | c.2945+1G>A | n/a | 0.0001 | 173226 | 33 | 0 | 0.0016 | 9.68E-01 | 38.79\|48.48\|n/a | 50.29\|51.21\|n/a | 49.06\|49.69\|n/a | 51.07\|52.65\|n/a | 4.42\|4.33\|n/a | 4.70\|5.00\|n/a | 4.25\|3.71\|n/a |
| chr6:160650337:C:T | rs533200556 | c.209+1G>A | n/a | 0.0001 | 173119 | 24 | 0 | 0.0008 | 9.77E-01 | 38.78\|66.67\|n/a | 50.29\|53.50\|n/a | 49.06\|52.56\|n/a | 51.07\|55.38\|n/a | 4.42\|4.41\|n/a | 4.70\|4.36\|n/a | 4.25\|4.50\|n/a |
| chr6:160650392:C:T | rs751841836 | c.155G>A | p.Trp52Ter | 0.0001 | 166587 | 29 | 0 | 0.0013 | 9.72E-01 | 39.23\|41.38\|n/a | 50.18\|51.48\|n/a | 48.97\|54.92\|n/a | 50.96\|49.06\|n/a | 4.44\|4.58\|n/a | 4.71\|5.00\|n/a | 4.27\|4.27\|n/a |
| chr6:160650438:G:A | rs121912503 | c.109C>T | p.Arg37Ter | 0.0001 | 173263 | 25 | 1 | 0.0011 | 0.00E+00 | 38.80\|40.00\|0.00 | 50.29\|47.36\|53.00 | 49.06\|51.60\|n/a | 51.07\|44.53\|53.00 | 4.42\|4.14\|4.00 | 4.70\|4.67\|n/a | 4.25\|3.75\|4.00 |
| chr6:160540042:C:A | n/a | c.5735+1G>T | n/a | 0 | 100032 | 2 | 0 | 0 | 9.97E-01 | 47.04\|50.00\|n/a | 47.58\|64.50\|n/a | 46.92\|65.00\|n/a | 48.17\|64.00\|n/a | 4.43\|4.50\|n/a | 4.65\|6.00\|n/a | 4.22\|3.00\|n/a |
| chr6:160540042:C:T | rs112462697 | c.5735+1G>A | n/a | 0 | 29023 | 1 | 0 | 0 | 9.98E-01 | 17.52\|0.00\|n/a | 53.82\|58.00\|n/a | 55.64\|n/a\|n/a | 53.43\|58.00\|n/a | 4.43\|8.00\|n/a | 4.97\|n/a\|n/a | 4.32\|8.00\|n/a |
| chr6:160540184:C:A | n/a | c.5595-1G>T | n/a | 0 | 29023 | 1 | 0 | 0 | 9.98E-01 | 17.52\|0.00\|n/a | 53.82\|55.00\|n/a | 55.64\|n/a\|n/a | 53.43\|55.00\|n/a | 4.43\|3.00\|n/a | 4.97\|n/a\|n/a | 4.32\|3.00\|n/a |
| chr6:160540185:T:C | n/a | c.5595-2A>G | n/a | 0 | 100002 | 5 | 0 | 0.0001 | 9.94E-01 | 47.04\|60.00\|n/a | 47.58\|40.20\|n/a | 46.92\|34.33\|n/a | 48.17\|49.00\|n/a | 4.43\|3.00\|n/a | 4.65\|0.00\|n/a | 4.22\|6.00\|n/a |
| chr6:160541106:C:T | n/a | c.5594+1G>A | n/a | 0 | 100038 | 1 | 0 | 0 | 9.99E-01 | 47.04\|0.00\|n/a | 47.58\|58.00\|n/a | 46.92\|n/a\|n/a | 48.17\|58.00\|n/a | 4.43\|2.00\|n/a | 4.65\|n/a\|n/a | 4.22\|2.00\|n/a |
| chr6:160541155:G:GT | n/a | c.5545dup | p.Thr1849AsnfsTer47 | 0 | 100030 | 1 | 0 | 0 | 9.99E-01 | 47.04\|0.00\|n/a | 47.58\|45.00\|n/a | 46.92\|n/a\|n/a | 48.17\|45.00\|n/a | 4.43\|3.00\|n/a | 4.65\|n/a\|n/a | 4.22\|3.00\|n/a |
| chr6:160542715:C:T | rs775800723 | c.5492G>A | p.Trp1831Ter | 0 | 137589 | 3 | 0 | 0 | 9.97E-01 | 43.81\|33.33\|n/a | 49.41\|56.67\|n/a | 48.41\|47.00\|n/a | 50.20\|61.50\|n/a | 4.45\|4.00\|n/a | 4.69\|3.00\|n/a | 4.25\|4.50\|n/a |
| chr6:160545469:AG:A | n/a | c.5368del | p.Leu1790PhefsTer32 | 0 | 166409 | 8 | 0 | 0.0001 | 9.92E-01 | 39.24\|25.00\|n/a | 50.18\|48.25\|n/a | 48.97\|38.00\|n/a | 50.96\|51.67\|n/a | 4.44\|3.33\|n/a | 4.71\|2.00\|n/a | 4.27\|4.00\|n/a |
| chr6:160545504:A:ACC | rs779555061 | c.5333_5334insGG | p.Asn1778LysfsTer9 | 0 | 29022 | 1 | 0 | 0 | 9.98E-01 | 17.52\|100.00\|n/a | 53.82\|60.00\|n/a | 55.64\|60.00\|n/a | 53.44\|n/a\|n/a | 4.43\|7.00\|n/a | 4.96\|7.00\|n/a | 4.32\|n/a\|n/a |
| chr6:160545534:C:T | n/a | c.5305-1G>A | n/a | 0 | 37445 | 1 | 0 | 0 | 9.98E-01 | 35.19\|0.00\|n/a | 54.27\|72.00\|n/a | 53.68\|n/a\|n/a | 54.59\|72.00\|n/a | 4.49\|8.00\|n/a | 4.83\|n/a\|n/a | 4.31\|8.00\|n/a |
| chr6:160547789:A:AT | rs754637067 | c.5303dup | p.Asn1768LysfsTer5 | 0 | 144117 | 4 | 0 | 0 | 9.96E-01 | 43.07\|75.00\|n/a | 49.58\|60.00\|n/a | 48.52\|56.67\|n/a | 50.38\|70.00\|n/a | 4.42\|4.50\|n/a | 4.67\|4.33\|n/a | 4.23\|5.00\|n/a |
| chr6:160547851:C:A | n/a | c.5242G>T | p.Glu1748Ter | 0 | 100038 | 1 | 0 | 0 | 9.99E-01 | 47.04\|100.00\|n/a | 47.58\|40.00\|n/a | 46.92\|40.00\|n/a | 48.17\|n/a\|n/a | 4.43\|7.00\|n/a | 4.65\|7.00\|n/a | 4.22\|n/a\|n/a |
| chr6:160547914:CT:C | rs1402894333 | c.5178del | p.Gly1727AspfsTer59 | 0 | 100029 | 2 | 0 | 0 | 9.97E-01 | 47.04\|0.00\|n/a | 47.58\|64.50\|n/a | 46.92\|n/a\|n/a | 48.18\|64.50\|n/a | 4.43\|7.50\|n/a | 4.65\|n/a\|n/a | 4.22\|7.50\|n/a |
| chr6:160548534:AC:A | n/a | c.5098del | p.Val1700TrpfsTer13 | 0 | 100038 | 1 | 0 | 0 | 9.99E-01 | 47.04\|100.00\|n/a | 47.58\|50.00\|n/a | 46.92\|50.00\|n/a | 48.17\|n/a\|n/a | 4.43\|7.00\|n/a | 4.65\|7.00\|n/a | 4.22\|n/a\|n/a |
| chr6:160548552:G:C | rs41267811 | c.5081C>G | p.Ser1694Ter | 0 | 29023 | 1 | 0 | 0 | 9.98E-01 | 17.52\|0.00\|n/a | 53.82\|48.00\|n/a | 55.64\|n/a\|n/a | 53.44\|48.00\|n/a | 4.43\|9.00\|n/a | 4.97\|n/a\|n/a | 4.32\|9.00\|n/a |
| chr6:160548559:G:A | rs757921434 | c.5074C>T | p.Arg1692Ter | 0 | 166607 | 7 | 0 | 0.0001 | 9.93E-01 | 39.23\|57.14\|n/a | 50.18\|53.00\|n/a | 48.97\|52.75\|n/a | 50.96\|53.33\|n/a | 4.44\|4.00\|n/a | 4.71\|3.50\|n/a | 4.27\|4.67\|n/a |
| chr6:160548611:AG:A | n/a | c.5021del | p.Pro1674LeufsTer17 | 0 | 29022 | 1 | 0 | 0 | 9.98E-01 | 17.52\|0.00\|n/a | 53.82\|61.00\|n/a | 55.64\|n/a\|n/a | 53.44\|61.00\|n/a | 4.43\|7.00\|n/a | 4.97\|n/a\|n/a | 4.32\|7.00\|n/a |
| chr6:160548638:G:T | n/a | c.4995C>A | p.Cys1665Ter | 0 | 137580 | 3 | 0 | 0 | 9.97E-01 | 43.81\|33.33\|n/a | 49.41\|64.00\|n/a | 48.41\|70.00\|n/a | 50.20\|61.00\|n/a | 4.45\|4.00\|n/a | 4.69\|1.00\|n/a | 4.25\|5.50\|n/a |
| chr6:160556023:A:T | n/a | c.4973+2T>A | n/a | 0 | 99191 | 1 | 0 | 0 | 9.99E-01 | 47.01\|100.00\|n/a | 47.58\|36.00\|n/a | 46.92\|36.00\|n/a | 48.16\|n/a\|n/a | 4.43\|8.00\|n/a | 4.65\|8.00\|n/a | 4.22\|n/a\|n/a |
| chr6:160556024:C:T | rs201888157 | c.4973+1G>A | n/a | 0 | 136122 | 9 | 0 | 0.0001 | 9.90E-01 | 43.80\|33.33\|n/a | 49.42\|47.44\|n/a | 48.41\|48.33\|n/a | 50.20\|47.00\|n/a | 4.45\|3.86\|n/a | 4.69\|6.50\|n/a | 4.25\|2.80\|n/a |
| chr6:160557442:T:TA | n/a | c.4760dup | p.Glu1588ArgfsTer16 | 0 | 100038 | 1 | 0 | 0 | 9.99E-01 | 47.04\|100.00\|n/a | 47.58\|35.00\|n/a | 46.92\|35.00\|n/a | 48.17\|n/a\|n/a | 4.43\|12.00\|n/a | 4.65\|12.00\|n/a | 4.22\|n/a\|n/a |
| chr6:160557448:AC:A | n/a | c.4754del | p.Gly1585ValfsTer3 | 0 | 100036 | 3 | 0 | 0 | 9.96E-01 | 47.04\|33.33\|n/a | 47.58\|50.67\|n/a | 46.92\|53.00\|n/a | 48.17\|49.50\|n/a | 4.43\|2.33\|n/a | 4.65\|3.00\|n/a | 4.22\|2.00\|n/a |
| chr6:160557491:CA:C | n/a | c.4711del | p.Trp1571GlyfsTer6 | 0 | 137586 | 3 | 0 | 0 | 9.97E-01 | 43.81\|33.33\|n/a | 49.41\|56.00\|n/a | 48.41\|55.00\|n/a | 50.20\|56.50\|n/a | 4.45\|4.00\|n/a | 4.69\|2.00\|n/a | 4.25\|5.00\|n/a |
| chr6:160557520:C:T | n/a | c.4683G>A | p.Trp1561Ter | 0 | 100038 | 1 | 0 | 0 | 9.99E-01 | 47.04\|0.00\|n/a | 47.58\|47.00\|n/a | 46.92\|n/a\|n/a | 48.17\|47.00\|n/a | 4.43\|3.00\|n/a | 4.65\|n/a\|n/a | 4.22\|3.00\|n/a |
| chr6:160557550:G:GCA | n/a | c.4651_4652dup | p.Arg1552AlafsTer19 | 0 | 100038 | 1 | 0 | 0 | 9.99E-01 | 47.04\|100.00\|n/a | 47.58\|48.00\|n/a | 46.92\|48.00\|n/a | 48.17\|n/a\|n/a | 4.43\|6.00\|n/a | 4.65\|6.00\|n/a | 4.22\|n/a\|n/a |
| chr6:160577134:A:G | rs1231300692 | c.4631+2T>C | n/a | 0 | 137460 | 10 | 0 | 0.0002 | 9.89E-01 | 43.80\|50.00\|n/a | 49.41\|51.40\|n/a | 48.41\|47.00\|n/a | 50.20\|55.80\|n/a | 4.45\|6.10\|n/a | 4.69\|5.40\|n/a | 4.25\|6.80\|n/a |
| chr6:160577135:C:A | n/a | c.4631+1G>T | n/a | 0 | 99788 | 1 | 0 | 0 | 9.99E-01 | 47.02\|100.00\|n/a | 47.58\|28.00\|n/a | 46.92\|28.00\|n/a | 48.17\|n/a\|n/a | 4.43\|1.00\|n/a | 4.65\|1.00\|n/a | 4.22\|n/a\|n/a |
| chr6:160578552:AC:A | n/a | c.4441del | p.Val1481PhefsTer49 | 0 | 29022 | 2 | 0 | 0 | 9.95E-01 | 17.52\|0.00\|n/a | 53.82\|56.50\|n/a | 55.64\|n/a\|n/a | 53.43\|56.50\|n/a | 4.43\|5.50\|n/a | 4.97\|n/a\|n/a | 4.32\|5.50\|n/a |
| chr6:160578604:G:A | rs1297292947 | c.4390C>T | p.Arg1464Ter | 0 | 166609 | 6 | 0 | 0.0001 | 9.94E-01 | 39.23\|33.33\|n/a | 50.18\|42.83\|n/a | 48.97\|41.50\|n/a | 50.96\|43.50\|n/a | 4.44\|5.33\|n/a | 4.71\|5.50\|n/a | 4.27\|5.25\|n/a |
| chr6:160578616:AG:A | n/a | c.4377del | p.Cys1460AlafsTer3 | 0 | 100038 | 1 | 0 | 0 | 9.99E-01 | 47.04\|0.00\|n/a | 47.58\|27.00\|n/a | 46.92\|n/a\|n/a | 48.17\|27.00\|n/a | 4.43\|4.00\|n/a | 4.65\|n/a\|n/a | 4.22\|4.00\|n/a |
| chr6:160578686:G:C | n/a | c.4308C>G | p.Tyr1436Ter | 0 | 100035 | 1 | 0 | 0 | 9.99E-01 | 47.04\|0.00\|n/a | 47.58\|63.00\|n/a | 46.92\|n/a\|n/a | 48.17\|63.00\|n/a | 4.43\|n/a\|n/a | 4.65\|n/a\|n/a | 4.22\|n/a\|n/a |
| chr6:160578705:C:T | n/a | c.4290-1G>A | n/a | 0 | 99997 | 1 | 0 | 0 | 9.99E-01 | 47.03\|0.00\|n/a | 47.58\|40.00\|n/a | 46.92\|n/a\|n/a | 48.18\|40.00\|n/a | 4.43\|5.00\|n/a | 4.65\|n/a\|n/a | 4.22\|5.00\|n/a |
| chr6:160578706:T:C | rs199583644 | c.4290-2A>G | n/a | 0 | 99967 | 0 | 1 | 0 | 0.00E+00 | 47.04\|n/a\|0.00 | 47.58\|n/a\|52.00 | 46.92\|n/a\|n/a | 48.18\|n/a\|52.00 | 4.43\|n/a\|8.00 | 4.65\|n/a\|n/a | 4.22\|n/a\|8.00 |
| chr6:160585045:C:A | rs41272114 | c.4289+1G>T | n/a | 0 | 100032 | 1 | 0 | 0 | 9.99E-01 | 47.04\|100.00\|n/a | 47.58\|25.00\|n/a | 46.92\|25.00\|n/a | 48.17\|n/a\|n/a | 4.43\|n/a\|n/a | 4.65\|n/a\|n/a | 4.22\|n/a\|n/a |
| chr6:160585099:C:T | rs1278172807 | c.4236G>A | p.Trp1412Ter | 0 | 100037 | 1 | 0 | 0 | 9.99E-01 | 47.04\|0.00\|n/a | 47.58\|67.00\|n/a | 46.92\|n/a\|n/a | 48.17\|67.00\|n/a | 4.43\|6.00\|n/a | 4.65\|n/a\|n/a | 4.22\|6.00\|n/a |
| chr6:160585107:G:A | n/a | c.4228C>T | p.Gln1410Ter | 0 | 37551 | 1 | 0 | 0 | 9.98E-01 | 35.21\|100.00\|n/a | 54.28\|55.00\|n/a | 53.70\|55.00\|n/a | 54.59\|n/a\|n/a | 4.49\|2.00\|n/a | 4.84\|2.00\|n/a | 4.31\|n/a\|n/a |
| chr6:160585146:G:A | rs200154828 | c.4189C>T | p.Arg1397Ter | 0 | 166603 | 11 | 0 | 0.0002 | 9.89E-01 | 39.23\|0.00\|n/a | 50.18\|55.36\|n/a | 48.97\|n/a\|n/a | 50.96\|55.36\|n/a | 4.44\|4.50\|n/a | 4.71\|n/a\|n/a | 4.27\|4.50\|n/a |
| chr6:160585158:C:A | n/a | c.4177G>T | p.Gly1393Ter | 0 | 35694 | 2 | 0 | 0 | 9.96E-01 | 19.48\|50.00\|n/a | 53.67\|60.00\|n/a | 54.70\|62.00\|n/a | 53.42\|58.00\|n/a | 4.35\|3.50\|n/a | 4.78\|2.00\|n/a | 4.25\|5.00\|n/a |
| chr6:160585167:G:A | rs146024102 | c.4168C>T | p.Arg1390Ter | 0 | 173285 | 3 | 0 | 0 | 9.97E-01 | 38.80\|33.33\|n/a | 50.29\|52.33\|n/a | 49.06\|44.00\|n/a | 51.07\|56.50\|n/a | 4.42\|3.00\|n/a | 4.70\|4.00\|n/a | 4.25\|2.50\|n/a |
| chr6:160585181:AC:A | n/a | c.4153del | p.Val1385SerfsTer31 | 0 | 100027 | 1 | 0 | 0 | 9.99E-01 | 47.04\|0.00\|n/a | 47.58\|71.00\|n/a | 46.92\|n/a\|n/a | 48.17\|71.00\|n/a | 4.43\|n/a\|n/a | 4.65\|n/a\|n/a | 4.22\|n/a\|n/a |
| chr6:160586532:GT:G | rs754513296 | c.4045del | p.Thr1349ArgfsTer5 | 0 | 166601 | 9 | 0 | 0.0001 | 9.91E-01 | 39.23\|55.56\|n/a | 50.18\|38.00\|n/a | 48.97\|34.20\|n/a | 50.96\|42.75\|n/a | 4.44\|3.50\|n/a | 4.71\|3.00\|n/a | 4.27\|4.00\|n/a |
| chr6:160586540:G:T | rs764320447 | c.4038C>A | p.Cys1346Ter | 0 | 166604 | 11 | 0 | 0.0002 | 9.89E-01 | 39.23\|54.55\|n/a | 50.18\|40.36\|n/a | 48.97\|35.50\|n/a | 50.96\|46.20\|n/a | 4.44\|3.56\|n/a | 4.71\|3.00\|n/a | 4.27\|4.00\|n/a |
| chr6:160589552:C:A | rs1032955724 | c.3947+1G>T | n/a | 0 | 29021 | 2 | 0 | 0 | 9.95E-01 | 17.52\|50.00\|n/a | 53.82\|65.50\|n/a | 55.63\|66.00\|n/a | 53.43\|65.00\|n/a | 4.43\|10.00\|n/a | 4.96\|10.00\|n/a | 4.32\|10.00\|n/a |
| chr6:160589581:G:A | rs1406394927 | c.3919C>T | p.Gln1307Ter | 0 | 37552 | 1 | 0 | 0 | 9.98E-01 | 35.22\|0.00\|n/a | 54.28\|58.00\|n/a | 53.70\|n/a\|n/a | 54.59\|58.00\|n/a | 4.49\|5.00\|n/a | 4.84\|n/a\|n/a | 4.31\|5.00\|n/a |
| chr6:160589585:C:T | rs780512843 | c.3915G>A | p.Trp1305Ter | 0 | 29022 | 1 | 0 | 0 | 9.98E-01 | 17.52\|100.00\|n/a | 53.82\|42.00\|n/a | 55.64\|42.00\|n/a | 53.43\|n/a\|n/a | 4.43\|5.00\|n/a | 4.97\|5.00\|n/a | 4.32\|n/a\|n/a |
| chr6:160589653:G:A | rs773141063 | c.3847C>T | p.Arg1283Ter | 0 | 135735 | 4 | 0 | 0 | 9.96E-01 | 39.79\|25.00\|n/a | 49.19\|45.75\|n/a | 47.92\|30.00\|n/a | 50.02\|51.00\|n/a | 4.41\|2.75\|n/a | 4.66\|4.00\|n/a | 4.23\|2.33\|n/a |
| chr6:160589657:ACT:A | rs758000323 | c.3841_3842del | p.Ser1281LeufsTer48 | 0 | 129061 | 1 | 0 | 0 | 9.99E-01 | 40.40\|100.00\|n/a | 48.99\|60.00\|n/a | 47.77\|60.00\|n/a | 49.81\|n/a\|n/a | 4.43\|5.00\|n/a | 4.68\|5.00\|n/a | 4.26\|n/a\|n/a |
| chr6:160590942:A:G | rs778727661 | c.3787+2T>C | n/a | 0 | 129049 | 2 | 0 | 0 | 9.98E-01 | 40.40\|0.00\|n/a | 48.99\|50.50\|n/a | 47.77\|n/a\|n/a | 49.81\|50.50\|n/a | 4.43\|3.00\|n/a | 4.68\|n/a\|n/a | 4.26\|3.00\|n/a |
| chr6:160590943:C:T | rs747762042 | c.3787+1G>A | n/a | 0 | 173278 | 12 | 0 | 0.0002 | 9.88E-01 | 38.80\|66.67\|n/a | 50.29\|48.50\|n/a | 49.06\|49.75\|n/a | 51.07\|46.00\|n/a | 4.42\|4.45\|n/a | 4.70\|4.86\|n/a | 4.25\|3.75\|n/a |
| chr6:160590965:AC:A | n/a | c.3765del | p.Ser1256ProfsTer46 | 0 | 144263 | 3 | 0 | 0 | 9.97E-01 | 43.08\|33.33\|n/a | 49.58\|65.67\|n/a | 48.52\|60.00\|n/a | 50.38\|68.50\|n/a | 4.42\|5.00\|n/a | 4.67\|4.00\|n/a | 4.23\|5.50\|n/a |
| chr6:160591001:G:A | rs1381325761 | c.3730C>T | p.Gln1244Ter | 0 | 100038 | 1 | 0 | 0 | 9.99E-01 | 47.04\|0.00\|n/a | 47.58\|26.00\|n/a | 46.92\|n/a\|n/a | 48.17\|26.00\|n/a | 4.43\|n/a\|n/a | 4.65\|n/a\|n/a | 4.22\|n/a\|n/a |
| chr6:160591050:C:T | rs1388217659 | c.3681G>A | p.Trp1227Ter | 0 | 66573 | 2 | 0 | 0 | 9.97E-01 | 27.50\|50.00\|n/a | 54.08\|64.00\|n/a | 54.24\|59.00\|n/a | 54.02\|69.00\|n/a | 4.46\|3.00\|n/a | 4.87\|4.00\|n/a | 4.31\|2.00\|n/a |
| chr6:160591098:GC:G | n/a | c.3632del | p.Gly1211AlafsTer2 | 0 | 29023 | 1 | 0 | 0 | 9.98E-01 | 17.52\|100.00\|n/a | 53.82\|65.00\|n/a | 55.63\|65.00\|n/a | 53.43\|n/a\|n/a | 4.43\|3.00\|n/a | 4.97\|3.00\|n/a | 4.32\|n/a\|n/a |
| chr6:160593986:G:A | n/a | c.3601C>T | p.Gln1201Ter | 0 | 37552 | 1 | 0 | 0 | 9.98E-01 | 35.22\|0.00\|n/a | 54.28\|53.00\|n/a | 53.70\|n/a\|n/a | 54.59\|53.00\|n/a | 4.49\|6.00\|n/a | 4.84\|n/a\|n/a | 4.31\|6.00\|n/a |
| chr6:160593990:C:T | n/a | c.3597G>A | p.Trp1199Ter | 0 | 100033 | 2 | 0 | 0 | 9.97E-01 | 47.04\|50.00\|n/a | 47.58\|50.00\|n/a | 46.92\|55.00\|n/a | 48.17\|45.00\|n/a | 4.43\|4.50\|n/a | 4.65\|8.00\|n/a | 4.22\|1.00\|n/a |
| chr6:160594011:C:T | n/a | c.3576G>A | p.Trp1192Ter | 0 | 37552 | 1 | 0 | 0 | 9.98E-01 | 35.22\|0.00\|n/a | 54.28\|62.00\|n/a | 53.70\|n/a\|n/a | 54.59\|62.00\|n/a | 4.49\|5.00\|n/a | 4.84\|n/a\|n/a | 4.31\|5.00\|n/a |
| chr6:160594020:A:T | n/a | c.3567T>A | p.Cys1189Ter | 0 | 37551 | 1 | 0 | 0 | 9.98E-01 | 35.21\|0.00\|n/a | 54.28\|51.00\|n/a | 53.70\|n/a\|n/a | 54.59\|51.00\|n/a | 4.49\|1.00\|n/a | 4.84\|n/a\|n/a | 4.31\|1.00\|n/a |
| chr6:160594029:TC:T | n/a | c.3557del | p.Gly1186GlufsTer10 | 0 | 29021 | 1 | 0 | 0 | 9.98E-01 | 17.52\|0.00\|n/a | 53.82\|54.00\|n/a | 55.64\|n/a\|n/a | 53.44\|54.00\|n/a | 4.43\|n/a\|n/a | 4.97\|n/a\|n/a | 4.32\|n/a\|n/a |
| chr6:160594080:G:T | n/a | c.3507C>A | p.Tyr1169Ter | 0 | 137583 | 8 | 0 | 0.0001 | 9.91E-01 | 43.81\|37.50\|n/a | 49.41\|59.75\|n/a | 48.41\|64.00\|n/a | 50.20\|57.20\|n/a | 4.45\|5.14\|n/a | 4.69\|6.00\|n/a | 4.25\|4.50\|n/a |
| chr6:160594102:CT:C | n/a | c.3484del | p.Ser1162AlafsTer34 | 0 | 99997 | 1 | 0 | 0 | 9.99E-01 | 47.03\|0.00\|n/a | 47.58\|23.00\|n/a | 46.92\|n/a\|n/a | 48.17\|23.00\|n/a | 4.43\|2.00\|n/a | 4.65\|n/a\|n/a | 4.22\|2.00\|n/a |
| chr6:160594106:G:A | n/a | c.3481C>T | p.Gln1161Ter | 0 | 100033 | 2 | 0 | 0 | 9.97E-01 | 47.04\|50.00\|n/a | 47.58\|50.50\|n/a | 46.92\|55.00\|n/a | 48.17\|46.00\|n/a | 4.43\|7.50\|n/a | 4.65\|7.00\|n/a | 4.22\|8.00\|n/a |
| chr6:160595353:C:A | rs1441650836 | c.3469+1G>T | n/a | 0 | 37552 | 1 | 0 | 0 | 9.98E-01 | 35.21\|100.00\|n/a | 54.28\|52.00\|n/a | 53.70\|52.00\|n/a | 54.59\|n/a\|n/a | 4.49\|1.00\|n/a | 4.84\|1.00\|n/a | 4.31\|n/a\|n/a |
| chr6:160595353:C:T | rs1441650836 | c.3469+1G>A | n/a | 0 | 129008 | 3 | 0 | 0 | 9.97E-01 | 40.39\|33.33\|n/a | 48.99\|56.33\|n/a | 47.77\|50.00\|n/a | 49.81\|59.50\|n/a | 4.43\|3.33\|n/a | 4.68\|5.00\|n/a | 4.26\|2.50\|n/a |
| chr6:160595368:GC:G | n/a | c.3454del | p.Ala1152LeufsTer44 | 0 | 99945 | 1 | 0 | 0 | 9.99E-01 | 47.03\|100.00\|n/a | 47.58\|55.00\|n/a | 46.92\|55.00\|n/a | 48.17\|n/a\|n/a | 4.43\|2.00\|n/a | 4.65\|2.00\|n/a | 4.22\|n/a\|n/a |
| chr6:160595485:C:T | rs752669457 | c.3338G>A | p.Trp1113Ter | 0 | 29023 | 1 | 0 | 0 | 9.98E-01 | 17.52\|0.00\|n/a | 53.82\|40.00\|n/a | 55.64\|n/a\|n/a | 53.44\|40.00\|n/a | 4.43\|6.00\|n/a | 4.97\|n/a\|n/a | 4.32\|6.00\|n/a |
| chr6:160599499:C:G | rs767906902 | c.3287+1G>C | n/a | 0 | 166451 | 3 | 0 | 0 | 9.97E-01 | 39.22\|66.67\|n/a | 50.18\|42.33\|n/a | 48.97\|45.00\|n/a | 50.96\|37.00\|n/a | 4.44\|1.00\|n/a | 4.71\|1.00\|n/a | 4.27\|1.00\|n/a |
| chr6:160599499:C:T | rs767906902 | c.3287+1G>A | n/a | 0 | 128898 | 2 | 0 | 0 | 9.98E-01 | 40.39\|50.00\|n/a | 48.99\|50.00\|n/a | 47.77\|60.00\|n/a | 49.81\|40.00\|n/a | 4.43\|3.50\|n/a | 4.68\|4.00\|n/a | 4.25\|3.00\|n/a |
| chr6:160599540:GT:G | n/a | c.3246del | p.Pro1083HisfsTer16 | 0 | 100027 | 1 | 0 | 0 | 9.99E-01 | 47.04\|0.00\|n/a | 47.58\|31.00\|n/a | 46.92\|n/a\|n/a | 48.17\|31.00\|n/a | 4.43\|2.00\|n/a | 4.65\|n/a\|n/a | 4.22\|2.00\|n/a |
| chr6:160599552:A:AC | n/a | c.3234dup | p.Ser1079ValfsTer9 | 0 | 100025 | 8 | 0 | 0.0002 | 9.90E-01 | 47.04\|62.50\|n/a | 47.58\|53.88\|n/a | 46.92\|56.40\|n/a | 48.17\|49.67\|n/a | 4.43\|3.29\|n/a | 4.65\|3.40\|n/a | 4.22\|3.00\|n/a |
| chr6:160599569:CTTCCTGT:C | rs773018558 | c.3211_3217del | p.Thr1071GlufsTer9 | 0 | 66564 | 2 | 0 | 0 | 9.97E-01 | 27.50\|100.00\|n/a | 54.08\|62.50\|n/a | 54.23\|62.50\|n/a | 54.02\|n/a\|n/a | 4.46\|8.00\|n/a | 4.87\|8.00\|n/a | 4.31\|n/a\|n/a |
| chr6:160599600:G:A | rs199952286 | c.3187C>T | p.Arg1063Ter | 0 | 100037 | 2 | 0 | 0 | 9.97E-01 | 47.04\|0.00\|n/a | 47.58\|51.00\|n/a | 46.92\|n/a\|n/a | 48.17\|51.00\|n/a | 4.43\|3.00\|n/a | 4.65\|n/a\|n/a | 4.22\|3.00\|n/a |
| chr6:160599622:G:T | rs756718229 | c.3165C>A | p.Tyr1055Ter | 0 | 100037 | 1 | 0 | 0 | 9.99E-01 | 47.04\|0.00\|n/a | 47.58\|60.00\|n/a | 46.92\|n/a\|n/a | 48.17\|60.00\|n/a | 4.43\|9.00\|n/a | 4.65\|n/a\|n/a | 4.22\|9.00\|n/a |
| chr6:160600916:C:T | n/a | c.3127+1G>A | n/a | 0 | 129006 | 8 | 0 | 0.0001 | 9.91E-01 | 40.40\|50.00\|n/a | 48.99\|56.88\|n/a | 47.77\|45.50\|n/a | 49.81\|68.25\|n/a | 4.43\|6.00\|n/a | 4.68\|6.25\|n/a | 4.25\|5.75\|n/a |
| chr6:160600923:C:CA | rs753812356 | c.3120dup | p.Glu1041Ter | 0 | 137207 | 11 | 1 | 0.0003 | 0.00E+00 | 43.80\|27.27\|0.00 | 49.42\|52.27\|35.00 | 48.41\|45.33\|n/a | 50.20\|54.88\|35.00 | 4.45\|4.44\|n/a | 4.69\|4.00\|n/a | 4.25\|4.67\|n/a |
| chr6:160600923:CA:C | rs753812356 | c.3120del | p.Phe1040LeufsTer5 | 0 | 99737 | 3 | 0 | 0 | 9.96E-01 | 47.01\|100.00\|n/a | 47.58\|48.00\|n/a | 46.91\|48.00\|n/a | 48.18\|n/a\|n/a | 4.43\|4.33\|n/a | 4.65\|4.33\|n/a | 4.22\|n/a\|n/a |
| chr6:160600998:G:A | rs142344792 | c.3046C>T | p.Arg1016Ter | 0 | 29022 | 1 | 0 | 0 | 9.98E-01 | 17.52\|0.00\|n/a | 53.82\|65.00\|n/a | 55.64\|n/a\|n/a | 53.43\|65.00\|n/a | 4.43\|6.00\|n/a | 4.97\|n/a\|n/a | 4.32\|6.00\|n/a |
| chr6:160601076:G:A | rs186072375 | c.2968C>T | p.Arg990Ter | 0 | 135728 | 9 | 0 | 0.0001 | 9.90E-01 | 39.79\|44.44\|n/a | 49.19\|44.56\|n/a | 47.92\|41.75\|n/a | 50.02\|46.80\|n/a | 4.40\|4.11\|n/a | 4.66\|2.50\|n/a | 4.23\|5.40\|n/a |
| chr6:160601080:G:T | n/a | c.2964C>A | p.Tyr988Ter | 0 | 100038 | 1 | 0 | 0 | 9.99E-01 | 47.04\|0.00\|n/a | 47.58\|35.00\|n/a | 46.92\|n/a\|n/a | 48.17\|35.00\|n/a | 4.43\|n/a\|n/a | 4.65\|n/a\|n/a | 4.22\|n/a\|n/a |
| chr6:160605045:C:A | n/a | c.2945+1G>T | n/a | 0 | 100007 | 1 | 0 | 0 | 9.99E-01 | 47.03\|100.00\|n/a | 47.58\|60.00\|n/a | 46.92\|60.00\|n/a | 48.17\|n/a\|n/a | 4.43\|3.00\|n/a | 4.65\|3.00\|n/a | 4.22\|n/a\|n/a |
| chr6:160605158:C:A | n/a | c.2833G>T | p.Gly945Ter | 0 | 100038 | 1 | 0 | 0 | 9.99E-01 | 47.04\|100.00\|n/a | 47.58\|56.00\|n/a | 46.92\|56.00\|n/a | 48.17\|n/a\|n/a | 4.43\|6.00\|n/a | 4.65\|6.00\|n/a | 4.22\|n/a\|n/a |
| chr6:160605181:AC:A | rs758029376 | c.2809del | p.Val937CysfsTer31 | 0 | 29022 | 1 | 0 | 0 | 9.98E-01 | 17.52\|100.00\|n/a | 53.82\|50.00\|n/a | 55.64\|50.00\|n/a | 53.43\|n/a\|n/a | 4.43\|5.00\|n/a | 4.97\|5.00\|n/a | 4.32\|n/a\|n/a |
| chr6:160605207:T:A | n/a | c.2786-2A>T | n/a | 0 | 37552 | 1 | 0 | 0 | 9.98E-01 | 35.21\|100.00\|n/a | 54.28\|36.00\|n/a | 53.70\|36.00\|n/a | 54.59\|n/a\|n/a | 4.49\|2.00\|n/a | 4.84\|2.00\|n/a | 4.31\|n/a\|n/a |
| chr6:160606476:C:T | rs1225171619 | c.2785+1G>A | n/a | 0 | 129008 | 2 | 0 | 0 | 9.98E-01 | 40.40\|0.00\|n/a | 48.99\|36.50\|n/a | 47.77\|n/a\|n/a | 49.81\|36.50\|n/a | 4.43\|5.00\|n/a | 4.68\|n/a\|n/a | 4.26\|5.00\|n/a |
| chr6:160606568:G:T | n/a | c.2694C>A | p.Cys898Ter | 0 | 37551 | 1 | 0 | 0 | 9.98E-01 | 35.22\|0.00\|n/a | 54.28\|50.00\|n/a | 53.70\|n/a\|n/a | 54.59\|50.00\|n/a | 4.49\|3.00\|n/a | 4.84\|n/a\|n/a | 4.31\|3.00\|n/a |
| chr6:160606587:CTG:C | rs1562342043 | c.2673_2674del | p.Ser892CysfsTer35 | 0 | 66567 | 3 | 0 | 0 | 9.95E-01 | 27.50\|0.00\|n/a | 54.08\|54.67\|n/a | 54.24\|n/a\|n/a | 54.02\|54.67\|n/a | 4.46\|4.33\|n/a | 4.87\|n/a\|n/a | 4.31\|4.33\|n/a |
| chr6:160606660:T:G | n/a | c.2604-2A>C | n/a | 0 | 100023 | 1 | 0 | 0 | 9.99E-01 | 47.04\|0.00\|n/a | 47.59\|60.00\|n/a | 46.92\|n/a\|n/a | 48.18\|60.00\|n/a | 4.43\|4.00\|n/a | 4.65\|n/a\|n/a | 4.22\|4.00\|n/a |
| chr6:160611615:C:T | n/a | c.2550G>A | p.Trp850Ter | 0 | 99722 | 1 | 0 | 0 | 9.99E-01 | 47.01\|0.00\|n/a | 47.59\|54.00\|n/a | 46.93\|n/a\|n/a | 48.18\|54.00\|n/a | 4.43\|2.00\|n/a | 4.65\|n/a\|n/a | 4.22\|2.00\|n/a |
| chr6:160611662:G:A | rs746733669 | c.2503C>T | p.Arg835Ter | 0 | 163751 | 6 | 0 | 0.0001 | 9.94E-01 | 39.14\|33.33\|n/a | 50.17\|54.33\|n/a | 48.95\|52.00\|n/a | 50.95\|55.50\|n/a | 4.44\|5.40\|n/a | 4.71\|2.00\|n/a | 4.27\|6.25\|n/a |
| chr6:160611684:G:T | rs774708289 | c.2481C>A | p.Tyr827Ter | 0 | 92602 | 1 | 0 | 0 | 9.99E-01 | 46.75\|100.00\|n/a | 47.59\|40.00\|n/a | 46.91\|40.00\|n/a | 48.19\|n/a\|n/a | 4.43\|5.00\|n/a | 4.65\|5.00\|n/a | 4.22\|n/a\|n/a |
| chr6:160635126:G:A | rs1406194254 | c.1072C>T | p.Gln358Ter | 0 | 29001 | 1 | 0 | 0 | 9.98E-01 | 17.52\|0.00\|n/a | 53.82\|78.00\|n/a | 55.64\|n/a\|n/a | 53.44\|78.00\|n/a | 4.43\|4.00\|n/a | 4.97\|n/a\|n/a | 4.32\|4.00\|n/a |
| chr6:160664164:A:G | rs1374200247 | c.49+2T>C | n/a | 0 | 133677 | 2 | 0 | 0 | 9.98E-01 | 40.10\|50.00\|n/a | 49.12\|43.50\|n/a | 47.88\|45.00\|n/a | 49.95\|42.00\|n/a | 4.40\|4.00\|n/a | 4.66\|3.00\|n/a | 4.22\|5.00\|n/a |
| chr6:160664201:TC:T | n/a | c.13del | p.Glu5LysfsTer10 | 0 | 99611 | 1 | 0 | 0 | 9.99E-01 | 47.02\|0.00\|n/a | 47.59\|62.00\|n/a | 46.92\|n/a\|n/a | 48.18\|62.00\|n/a | 4.43\|n/a\|n/a | 4.65\|n/a\|n/a | 4.22\|n/a\|n/a |

### Analysis of *LPA* loss-of-function (LoF) in the Pakistan Genome Resource

For each *LPA* LoF variant in PGR, we compared expected homozygotes under Hardy-Weinburg equilibrium with observed counts (**Table 1**). No variant showed significant homozygote depletion, suggesting no embryonic lethality or major reproductive fitness effect, though high consanguinity may mask subtle effects.

Consanguinity led to homozygote enrichment, notably for c.4289+1G>A. A total of 404 homozygote males and females were observed, compared to 233 expected based on the allele frequencies observed in PGR and the assumptions of Hardy Weinberg Equilibrium. Females compromised 38% of LoF homozygotes, consistent with the overall percentage of female representation across the entire PGR cohort (39%). Male over representation reflects cardiovascular disease-focused recruitment.

Child count data were available for most participants. Mean numbers of children by zygosity and sex are shown in **Table 1**. Burden analysis tested associations with ever having children (binary) and total number (quantitative) (**Table 2**). No significant associations were found when data were aggregated by sex or when specifically evaluated for a particular sex.

**Table 2.** PGR *LPA* LoF burden association statistics. *LPA* LoF burden association stats for number of children and having had any children under additive and recessive models for all of PGR, females in PGR, and males in PGR.

| Phenotype | Subset | Additive Beta [95% CI] | Recessive Beta [95% CI] | Additive OR [95% CI] | Recessive OR [95% CI] | Additive P-value | Recessive P-value |
|---|---|---|---|---|---|---|---|
| Has children | All | 0.03<br>[-0.06, 0.13] | -0.25<br>[-0.75, 0.25] | 1.03<br>[0.94, 1.14] | 0.78<br>[0.47, 1.28] | 0.52 | 0.32 |
| Has children | Female | -0.01<br>[-0.18, 0.17] | -0.83<br>[-1.75, 0.09] | 0.99<br>[0.83, 1.18] | 0.44<br>[0.17, 1.10] | 0.93 | 0.08 |
| Has children | Male | 0.08<br>[-0.04, 0.20] | 0.05<br>[-0.60, 0.71] | 1.08<br>[0.96, 1.22] | 1.06<br>[0.55, 2.04] | 0.19 | 0.87 |
| Number of children | All | -0.01<br>[-0.02, 0.01] | -0.03<br>[-0.13, 0.06] | n/a | n/a | 0.52 | 0.45 |
| Number of children | Female | 0.00<br>[-0.03, 0.03] | -0.09<br>[-0.24, 0.07] | n/a | n/a | 0.90 | 0.27 |
| Number of children | Male | 0.00<br>[-0.02, 0.03] | 0.03<br>[-0.08, 0.15] | n/a | n/a | 0.66 | 0.58 |

### Recall and reproductive health history characterization of *LPA* loss-of-function (LoF) female homozygotes and controls

To better understand whether lifelong deficiency of Lp(a) due to homozygous germline LoF variants in the *LPA* gene was associated with self-reported reproductive effects, women participants from PGR were recontacted to profile reproductive history. A total of 157 *LPA* LoF female homozygote and 157 female matched controls were contacted by phone for participation in a study that would query aspects of their reproductive history and the health of their children (see **Figure 2** for a CONSORT flow diagram). Among these, 13 females homozygous LoF and 13 non-carrier female controls participated. Sanger sequencing of these probands and their consenting family members identified a total of 18 LoF homozygous females, 7 LoF homozygous males, 54 LoF heterozygous females, 43 LoF heterozygous males, 133 homozygous reference (non-carrier) females, and 106 homozygous reference males. Additionally, one female was heterozygous for two *LPA* LoF alleles (rs41272114 and rs559580002) and was treated as a compound heterozygote and grouped with the *LPA* LoF homozygotes for all subsequent analyses. Participant demographics are shown in **Table 3**. Overall, 19 homozygous LoF females (including one presumed compound heterozygote), 31 heterozygous LoF females, and 86 homozygous reference females were surveyed for their reproductive health and pregnancy outcomes.

**Figure 2.**
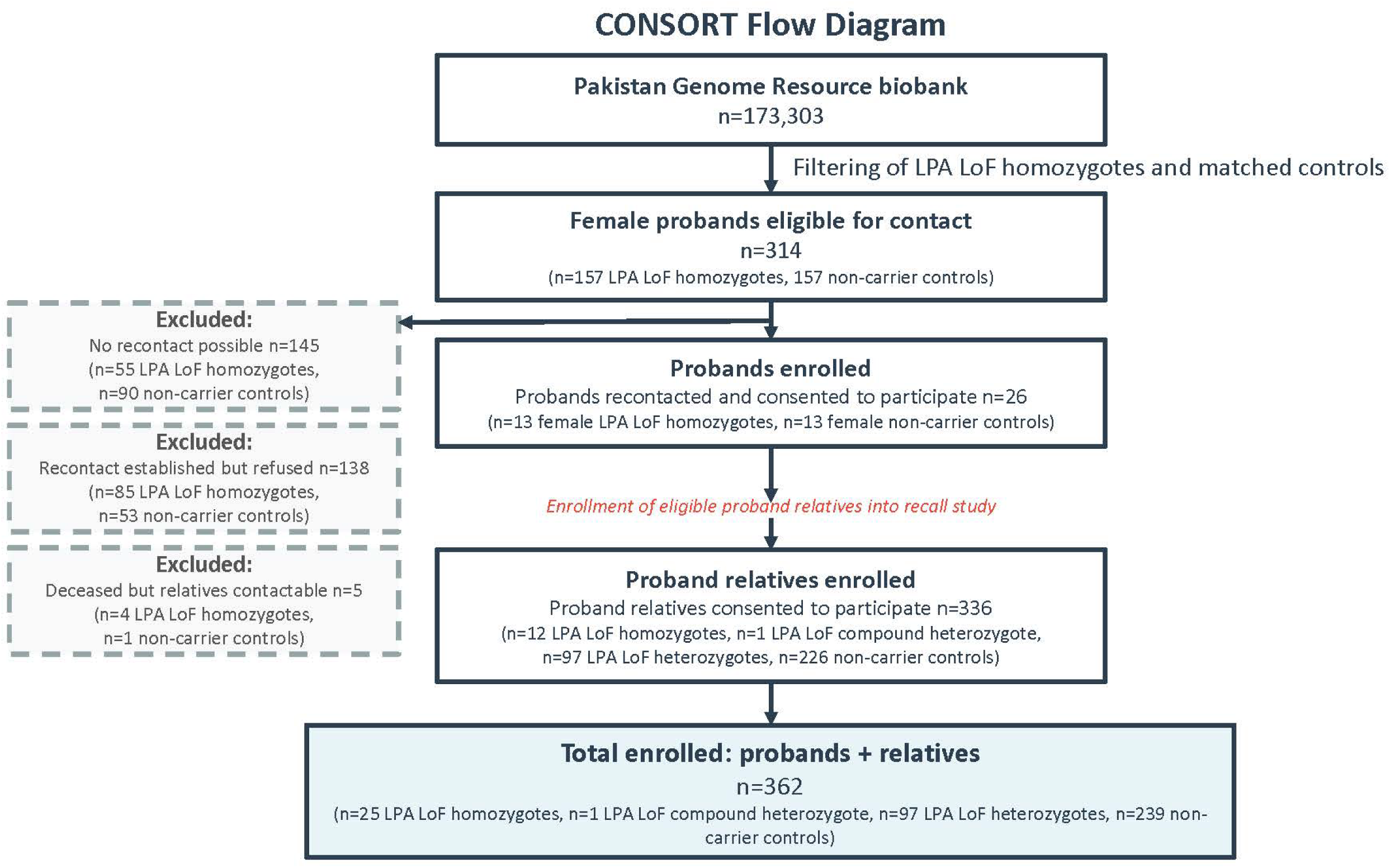
CONSORT flow diagram. Diagram showing the number of participants included in the recall by genotype study by LPA LoF genotypes

**Table 3.** Recall by genotype study sample summary statistics. Demographic breakdown of all recall participants (top) as well as those female recall participants surveyed for their pregnancy outcomes (bottom).

| **All Recall Study Participants** |  |  |  |  |  |  |  |  |  |
|---|---|---|---|---|---|---|---|---|---|
|  | **All Individuals** |  |  | **Males** |  |  | **Females** |  |  |
| **Phenotype** | **Homozygous Reference** | **Heterozygous LoF** | **Homozygous LoF** | **Homozygous Reference** | **Heterozygous LoF** | **Homozygous LoF** | **Homozygous Reference** | **Heterozygous LoF** | **Homozygous LoF** |
| N | 239 | 97 | 26 | 106 | 43 | 7 | 133 | 54 | 19 |
| Female (%) | 133 (55.6%) | 54 (55.7%) | 19 (73.1%) | N/A | N/A | N/A | 133 (100.0%) | 54 (100.0%) | 19 (100.0%) |
| Age (years) | 33.2 (16.4) | 30.6 (16.1) | 46.6 (18.2) | 34.1 (17.8) | 32.0 (18.4) | 37.0 (24.4) | 32.5 (15.2) | 29.6 (14.1) | 50.2 (14.6) |
| BMI (kg/m<sup>2</sup>) | 23.4 (5.9) | 23.1 (5.0) | 25.7 (6.2) | 23.4 (6.1) | 22.5 (3.9) | 22.3 (2.9) | 23.4 (5.8) | 23.6 (5.7) | 26.7 (6.7) |
| Number of children | 2.4 (2.6) | 1.9 (2.6) | 4.1 (3.5) | 2.3 (2.7) | 1.6 (2.5) | 2.7 (4.4) | 2.5 (2.5) | 2.2 (2.6) | 4.6 (3.2) |
| Hypertension | 41 (17.2%) | 16 (16.5%) | 12 (46.2%) | 17 (16.0%) | 4 (9.3%) | 1 (14.3%) | 24 (18.0%) | 12 (22.2%) | 11 (57.9%) |
| Diabetes | 27 (11.3%) | 10 (10.3%) | 8 (30.8%) | 12 (11.3%) | 3 (7.0%) | 1 (14.3%) | 15 (11.3%) | 7 (13.0%) | 7 (36.8%) |
| Pre-eclampsia | 8 (3.3%) | 8 (8.2%) | 1 (3.8%) | N/A | N/A | N/A | 8 (6.0%) | 8 (14.8%) | 1 (5.3%) |
| **Study Participants Surveyed for Pregnancy Outcomes** |  |  |  |  |  |  |  |  |  |
| **Phenotype** | **Homozygous Reference** | **Heterozygous LoF** | **Homozygous LoF** |  |  |  |  |  |  |
| N | 86 | 31 | 19 |  |  |  |  |  |  |
| Female (%) | 86 (100.0%) | 31 (100.0%) | 19 (100.0%) |  |  |  |  |  |  |
| Age (years) | 39.3 (13.6) | 37.5 (12.6) | 50.2 (14.6) |  |  |  |  |  |  |
| BMI (kg/m<sup>2</sup>) | 24.9 (5.3) | 25.9 (6.0) | 26.7 (6.7) |  |  |  |  |  |  |
| Number of children | 3.7 (2.1) | 3.7 (2.5) | 4.6 (3.2) |  |  |  |  |  |  |
| Hypertension | 22 (25.6%) | 12 (38.7%) | 11 (57.9%) |  |  |  |  |  |  |
| Diabetes | 15 (17.4%) | 7 (22.6%) | 7 (36.8%) |  |  |  |  |  |  |
| Pre-eclampsia | 8 (9.3%) | 8 (25.8%) | 1 (5.3%) |  |  |  |  |  |  |

#### Pregnancy and delivery history

The recall history survey identified 107 pregnancies in 19 LoF homozygous women (including the compound heterozygote), 141 pregnancies in 31 LoF heterozygous women, and 401 in 86 homozygous reference women (**Table 4**; **Figure 3a**).

**Figure 3.**
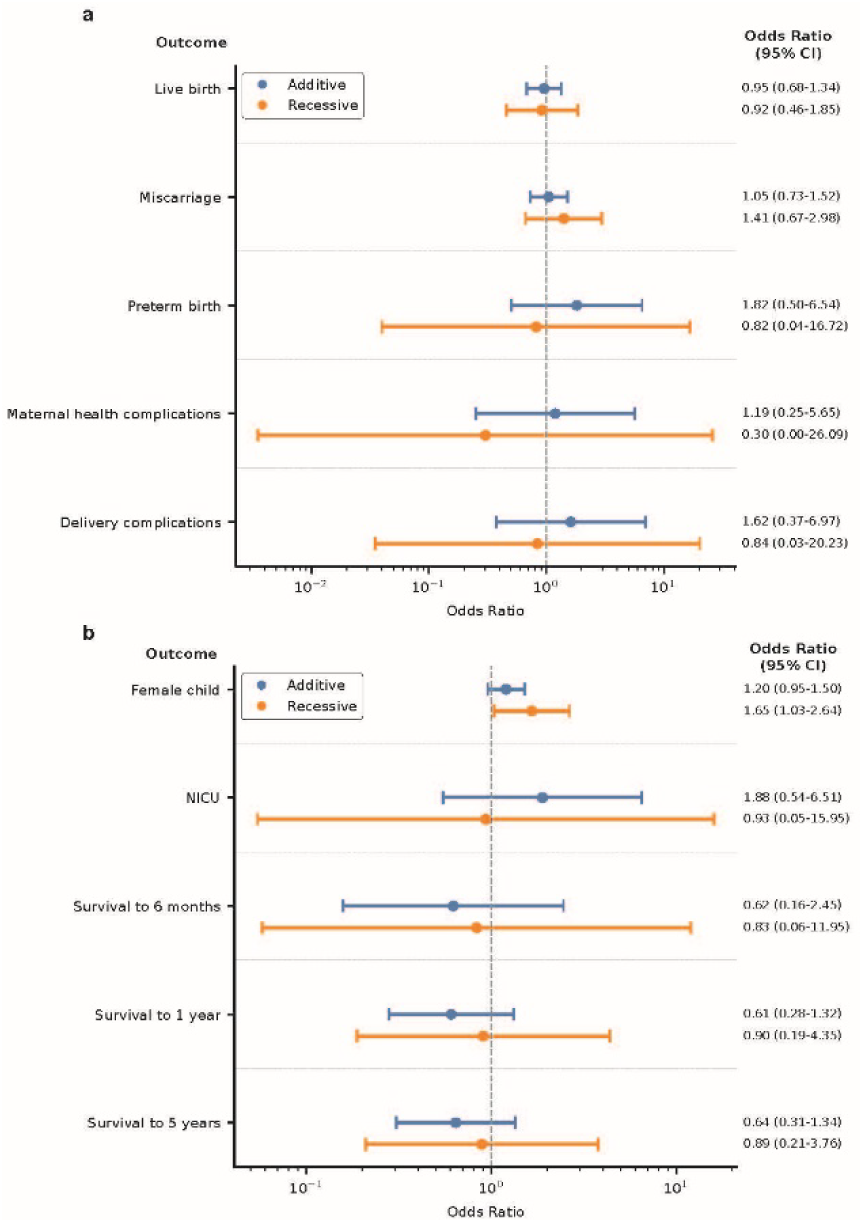
Pregnancy outcomes in recall by genotype study. Association statistics for pregnancy-related outcomes **(a)** and perinatal and post-natal outcomes **(b)**. Results for birth defects are included in the main text but excluded in the figure due to wide confidence intervals. Points represent odds ratios; horizontal bars represent odds ratio 95% confidence intervals. Odds ratios are plotted on a log-scale. NICU, neonatal intensive care unit; CI, confidence interval.

**Table 4.** Recall by genotype study pregnancy outcomes. Number of occurrences of various pregnancy outcomes by genotype.

| LPA LoF Genotype | Women Surveyed | Pregnancies | Sets of Twins | Miscarriages | Still Births | Total Abortions | Medical Abortions | Non-medical Abortions | Pregnancies with a Live Birth | Preterm Births | Pregnancies with Maternal Complications | Deliveries with Maternal Complications |
|---|---|---|---|---|---|---|---|---|---|---|---|---|
| Homozygous Reference | 86 | 401 | 7 (1.7%) | 69 (17.2%) | 2 (0.5%) | 11 (2.7%) | 5 (1.2%) | 6 (1.5%) | 314 (78.3%) | 6 (1.5%) | 15 (3.7%) | 36 (9.0%) |
| Heterozygous LoF | 31 | 141 | 2 (1.4%) | 19 (13.5%) | 7 (5.0%) | 3 (2.1%) | 1 (0.7%) | 2 (1.4%) | 112 (79.4%) | 10 (7.1%) | 25 (17.8%) | 33 (23.4%) |
| Homozygous LoF | 19 | 107 | 3 (2.8%) | 21 (19.6%) | 1 (0.9%) | 0 (0.0%) | 0 (0.0%) | 0 (0.0%) | 84 (78.5%) | 3 (2.8%) | 2 (1.9%) | 9 (8.4%) |

Number of pregnancies did not differ by *LPA* LoF genotype (β_additive_=-0.09 pregnancies, 95% CI [-0.62, 0.45], P=0.75; β_recessive_=-0.31 [-1.45, 0.84], P=0.60). All 19 female LoF homozygous women had at least one pregnancy. Three LoF homozygote (15.8% of women and 3.6% of pregnancies with a live birth), two LoF heterozygote (6.5% of women and 1.8% of pregnancies with a live birth), and 7 homozygous reference individuals (8.1% of women and 2.2% of pregnancies with a live birth) each reported having one set of twins. *LP*A LoF genotype was not associated with the odds of a pregnancy resulting in a livebirth (OR_additive_=0.95, [0.68, 1.34], P_additive_=0.67; OR_recessive_=0.92, [0.46, 1.85], P_recessive_=0.81), or miscarriage (OR_additive_=1.05 [0.73, 1.52], P=0.78; OR_recessive_=1.41 [0.67, 2.98], P=0.37).

Pregnancy-loss event rates (stillbirth, miscarriage, abortion) were comparable across groups. One stillbirth (0.9% of pregnancies), 21 miscarriages (19.6% of pregnancies), and 0 abortions were reported by *LPA* LoF homozygotes. 7 stillbirths (5.0% of pregnancies), 19 miscarriages (13.4% of pregnancies), and 3 abortions (2.1% of pregnancies; 1 for a medical reason and 2 for non-medical personal reasons) were reported by *LPA* LoF heterozygotes. 2 stillbirths (0.5% of pregnancies), 69 miscarriages (17.2% of pregnancies), and 11 abortions (2.7% of pregnancies; 5 for medical reasons and 6 for non-medical personal reasons) were reported by *LPA* homozygous reference individuals.

Preterm birth was not associated with LoF genotype (OR_additive_=1.82 [0.50, 6.54], P=0.36; OR_recessive_=0.82 [0.04,16.72], P=0.90) with similar rates across groups. Three pre-term births were reported by *LPA* LoF homozygotes (2.8% of pregnancies). 10 preterm births were reported by *LPA* LoF heterozygotes (7.1% of pregnancies). 6 preterm births were reported by *LPA* homozygous reference individuals (1.5% of pregnancies).

Maternal-health complications were not associated with LoF genotype (OR_additive_=1.19, [0.25, 5.62], P_additive_=0.83; OR_recessive_=0.30, [0.00, 26.09], P_recessive_=0.60) with similar distributions of hypertension, gestational diabetes, and preeclampsia. Two *LPA* LoF homozygotes (10.5%) reported complications with one of their pregnancies. Specifically, one woman reported hypertension (which may include gestational hypertension or chronic hypertension), and one woman reported both pre-eclampsia and gestational diabetes. 10 *LPA* LoF heterozygotes (32.2%) reported complications with at least one pregnancy, including hypertension (6 women), gestational diabetes (3 women), and preeclampsia (3 women). 12 *LPA* homozygous reference women (14.0%) reported complications with at least one pregnancy, including preeclampsia (8 women), hypertension (2 women), and gestational diabetes (2 women).

Delivery complications were also not associated with LoF genotype (OR_additive_=1.62, [0.37, 6.97], P_additive_=0.52; OR_recessive_=0.84, [0.03, 20.23], P_recessive_=0.91), with broadly similar complication patterns. Five *LPA* LoF homozygotes (26.3%) reported complications with at least one delivery, including C-sections (2 women), hypertension (1 woman), anemia (1 woman), and undisclosed complications (1 woman). 16 *LPA* LoF heterozygotes (51.6%) reported complications with at least one delivery, including C-sections (10 women), hypertension (4 women), preeclampsia (3 women), fetal macrosomia (1 woman), distress to the baby (1 woman), induced labor (1 woman), and undisclosed complications (1 woman). 17 *LPA* homozygous reference women (19.8%) reported complications with at least one pregnancy, including C-sections (9 women), preeclampsia (1 woman), induced labor (3 women), leakage of amniotic fluid (1 woman), low fetal heart rate (1 woman), and undisclosed complications (2 women).

#### Perinatal and postnatal outcomes

Perinatal and postnatal outcomes are summarized in **Table 5** and **Figure 3b**. Number of children did not differ by LoF genotype (β_additive_=-0.09 children, [-0.54, 0.36], P=0.69; β_recessive_=-0.35, [-1.31, 0.60], P=0.46). 87 children were born to LoF homozygote women (4.6 children per woman), 114 children were born to LoF heterozygotes (3.7 children per woman), and 321 to homozygous reference women (3.7 children per woman). Most LoF homozygote women (18/19) had at least one child, with 16 having multiple children.

**Table 5.**
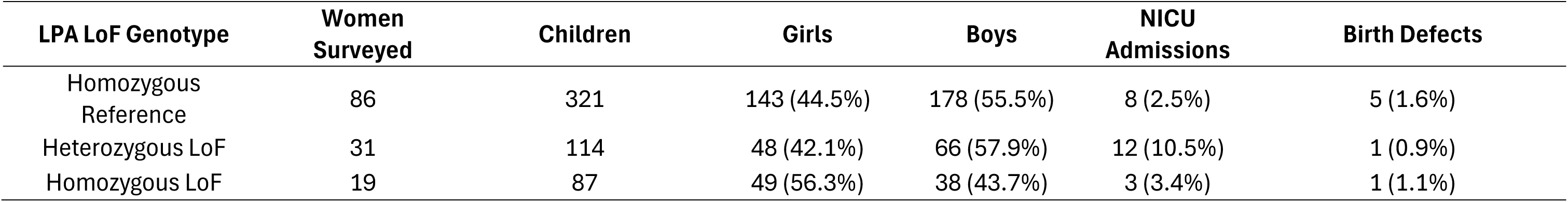
Recall by genotype study child outcomes. Number of occurrences of various child outcomes by genotype.

| **LPA LoF Genotype** | **Women Surveyed** | **Children** | **Girls** | **Boys** | **NICU Admissions** | **Birth Defects** |
|---|---|---|---|---|---|---|
| Homozygous Reference | 86 | 321 | 143 (44.5%) | 178 (55.5%) | 8 (2.5%) | 5 (1.6%) |
| Heterozygous LoF | 31 | 114 | 48 (42.1%) | 66 (57.9%) | 12 (10.5%) | 1 (0.9%) |
| Homozygous LoF | 19 | 87 | 49 (56.3%) | 38 (43.7%) | 3 (3.4%) | 1 (1.1%) |

Female sex was nominally more likely under a recessive model (OR_additive_=1.20 [0.95, 1.50], P=0.12; OR_recessive_=1.65 [1.04, 2.64], P=0.04). 56% of offspring of LoF homozygotes were female.

Neonatal intensive care unit admission was not associated with *LPA* LoF genotype (OR_additive_=1.88 [0.54, 6.51], P=0.32; OR_recessive_=0.93 [0.05, 15.95], P=0.96) with similar counts across groups. Three children of LoF homozygotes (3.4% of children) were admitted to the neonatal intensive care unit, compared to 12 children (10.5%) of LoF heterozygotes, and 8 children (2.5%) of homozygous reference individuals.

Birth defects were not associated with LoF genotype (OR_additive_=0.74 [0.01, 37.09], P=0.88; OR_recessive_=0.74 [0.00, 2018.74], P=0.94) and were rare across groups. One child of an LoF homozygote was born with a birth defect (neurological disorder), as compared to 1 child of a LoF heterozygote (encephalopathy), and 3 children of homozygous reference women (reported as stomach problem, missing kidney, and undisclosed reason).

Child survival and health at 6 months, 1 year, and 5 years were similar across genotypes (**Table 6**).

**Table 6.** Status of children at 6 months, 1 year, and 5 years. The percentage of children reported as healthy or unhealthy at each age is relative to the total number of children having reached that age. The percentage of children reported to have died between 6 months to 1 year is relative to the total number of children having survived to 6 months. Similarly, the percentage of children reported to have died between 1 year and 5 years is relative to the total number of children having survived to 1 year.

| **Maternal LPA LoF Genotype** | **Died before 6 months** | **Healthy at 6 months** | **Unhealthy at 6 months** | **Died 6 months to 1 year** | **Healthy at 1 year** | **Unhealthy at 1 year** | **Died 1 year to 5 years** | **Healthy at 5 years** | **Unhealthy at 5 years** |
|---|---|---|---|---|---|---|---|---|---|
| Homozygous Reference | 8 (2.8%) | 268 (97.5%) | 7 (2.5%) | 3 (1.1%) | 264 (98.5%) | 4 (1.5%) | 2 (0.8%) | 232 (98.7%) | 3 (1.3%) |
| Heterozygous LoF | 8 (7.3%) | 98 (96.1%) | 4 (3.9%) | 3 ( 3.0%) | 94 (96.9%) | 3 (3.1%) | 0 (0.0%) | 81 (97.6%) | 2 (2.4%) |
| Homozygous LoF | 3 (3.9%) | 70 (94.6%) | 4 (5.4%) | 2 (2.7%) | 70 (97.2%) | 2 (2.8%) | 1 (1.4%) | 65 (95.6%) | 3 (4.4%) |

#### 6 months

The odds of a child surviving to 6 months were not associated with maternal *LPA* LoF genotype (OR_additive_= 0.62 [0.16, 2.45], P=0.50; OR_recessive_= 0.83 [0.06, 11.95], P=0.89). Survival was 93–97% across groups, with few illnesses. 74 out of 77 children (96%) of LoF homozygotes lived to the age of 6 months, out of which 4 were reported unhealthy (2 due to polio, 2 due to chickenpox, and 2 for undisclosed reasons). 102 out of 110 children (93%) of LoF heterozygotes lived to the age of 6 months, out of which 4 were reported unhealthy (2 due to polio and 2 for undisclosed reasons). 275 out of 283 children (97%) of homozygous reference individuals lived to the age of 6 months, out of which 7 were reported unhealthy (2 due to polio, 2 due to respiratory problems, and 3 for undisclosed reasons).

#### 1 year

The odds of a child surviving to 1 year were not associated with maternal LoF genotype (OR_additive_= 0.61 [0.28, 1.32], P=0.21; OR_recessive_= 0.90 [0.19, 4.35], P=0.90). 72 out of 77 children (94%) of LoF homozygotes lived to the age of 1 year, out of which 2 were reported unhealthy (1 due to fits and 1 for undisclosed reasons). 97 out of 108 children (90%) of LoF heterozygotes lived to the age of 1 year, out of which 3 were reported unhealthy (2 due to polio and 1 for undisclosed reasons). 268 out of 279 children (96%) of homozygous reference individuals lived to the age of 1 year, out of which 4 were reported unhealthy (2 due to respiratory problems, 1 due to recurrent chest infection and mumps, and 1 due to chickenpox).

#### 5 years

The odds of a child surviving to 5 years were not associated with maternal LoF genotype (OR_additive_= 0.64 [0.31, 1.34], P=0.23; OR_recessive_= 0.89 [0.21, 3.76], P=0.87). Reported illnesses were infrequent and varied, with no consistent genotype pattern. 68 out of 74 children (92%) of LoF homozygotes lived to the age of 5 years, out of which 3 were reported unhealthy (1 due to fits and 2 for undisclosed reasons). 83 out of 94 children (88%) of LoF heterozygotes lived to the age of 5 years, out of which 2 were reported unhealthy (both due polio). 235 out of 248 children (95%) of homozygous reference individuals lived to the age of 5 years, out of which 3 were reported unhealthy (1 due to chickenpox, 1 due to tuberculosis, and 1 for undisclosed reasons).

## Discussion

This study evaluates whether lifelong Lp(a) deficiency due to homozygous *LPA* LoF variants affects reproductive fitness or maternal–child health, informing safety of Lp(a)-lowering therapies. Prior human data on reproductive traits are limited. Early reports described two small families with LoF splice variants, including one homozygous null woman with a heterozygous daughter^27^. In Finland, 227 homozygous/compound heterozygous individuals showed no increased morbidity or mortality^28^. Although the genotypes of the parents of these participants are not known, the prevalence of heterozygous and homozygous carriers of these variants was found to be in Hardy-Weinberg equilibrium (HWE). This study expands prior work^15, 32^ by including a substantially higher number of cases with detailed evaluation of maternal fetal pairs and reproductive history.

Using the PGR, a large, consanguineous South Asian biobank, we further assessed reproductive traits amongst individuals with *LPA* LoF variants. Multiple high-confidence *LPA* LoF variants were identified that markedly reduce circulating Lp(a) in a gene-dosage–dependent manner. Variants showed no depletion of homozygotes vs HWE and no association with fertility or number of children. These findings suggest no overt population-level effect on reproductive fitness.

In a genotype-based recall study, 19 female *LPA* LoF homozygotes and controls were surveyed on pregnancy, delivery, and offspring outcomes. Across 107 pregnancies and 87 children in homozygous women, rates of pregnancies, live births, miscarriages, maternal complications (including preeclampsia, hypertension, gestational diabetes), delivery complications, neonatal intensive care admissions, birth defects, and child health problems were comparable to those in heterozygous and non-carrier women, with no statistically significant genotype–outcome associations.

Furthermore, this study demonstrates the utility of using rare genetic variants to understand on-target perturbations of specific therapeutic proteins or pathways when these differ in substantial ways between humans and preclinical species. In this case rodents do not express apolipoprotein(a) or Lp(a), and non-human primates have sequence variance in the plasminogen like domain such that functional studies could miss relevant safety outcomes. In this case, human reproductive data best informs potential safety concerns, of which none were identified, for women who would benefit from targeted Lp(a) lowering. This approach could be more broadly applied.

## Limitations

This study has limitations. First, while the total number of LPA LoF individuals identified further confirmed HWE principals, recall participation rate was low (8%) among recontacted probands raising potential selection bias and the direction of this bias cannot be determined, limiting generalizability. However, detailed reproductive history was previously limited to two small families^27^. Second, despite age matching prior to recontact, *LPA* LoF homozygotes were older on average than heterozygotes and non-carriers (**Table 3**), as family members of probands tended to be younger individuals who were disproportionately non-carriers or heterozygotes. Third, outcomes were self-reported introducing recall and comprehension bias, particularly for events such as early miscarriage. Finally, rare events (*e.g*., stillbirth) yielded wide confidence intervals—especially for recessive models with few homozygotes— limiting power to detect modest effects. Finally, genetic LoF individuals have lifelong low Lp(a) and may not reflect reproductive findings in those with high Lp(a) that is reduced therapeutically.

## Conclusion

Overall, our findings demonstrate complete lifelong *LPA* deficiency is not associated with evident adverse maternal reproductive outcomes or major perinatal/postnatal child health issues, supporting the reproductive safety of potent Lp(a)-lowering therapies that mimic genetic loss of function.

## Acknowledgements

We are grateful to the Pakastani participants of the CNDC who contributed to the work, without whom this would not have been possible.

## Supplement

### Primer design

Primers were designed for the 3 variants targeted for recall by genotype studies using Primer 3Plus Software, and *in-silico* PCR was run using the UCSC Genome Browser’s tool. **Supplemental Table 1** shows primer sequences.

**Supplemental Table 1.**
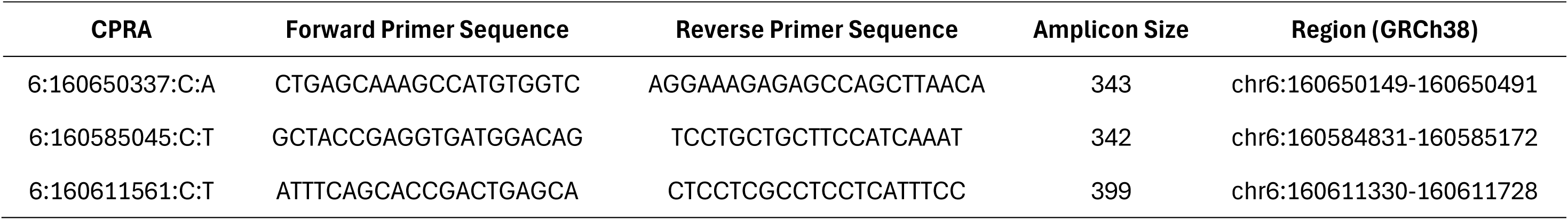
Primer sequences used for Sanger sequencing. Primer sequences used for Sanger sequencing of 3 most common *LPA* LoF variants.

### General assessments for the recall by genotype

Physical anthropometric information was collected including height, weight, waist circumference, and hip circumference that were taken in standing position by using height and weight scales and a spring gauge tape measure. Blood pressure and heart rate were recorded by using OMRON healthcare M2 blood pressure monitors. Non-fasting blood samples were collected and processed from each participant. BD Vacutainer SST tubes (cat# 367986) were used for sera collection and BD Vacutainer EDTA tubes (cat# 367856) for plasma collection. Serum and plasma were separated within 45 minutes of venipuncture. A random urine sample was also collected from each participant. The samples were stored temporarily in dry ice in the field and transported to a central laboratory based at CNCD and stored at -80 degrees Celsius. Measurements for total-cholesterol, HDL cholesterol, LDL cholesterol, triglycerides, VLDL, AST, ALT and creatinine were made in serum samples using enzymatic assays, whereas concentrations of HbA1c were measured using a turbidimetric assay in whole-blood samples (Roche Diagnostics, USA). Urinary microalbumin was also measured through the urine sample.

### Recall by genotype: Survey related to reproductive health were

1. How many times have you been pregnant?
2. How many children have you had?

◦ How many live births?
◦ How many still births?
◦ How many multiples (twins/triples/others)?
3. Did you have any miscarriage(s)?

◦ How many?
4. Did you have any abortion(s)?

◦ Were any for medical reasons? If so, how many?
◦ Were any for non-medical or personal reasons? If so, how many?
5. Did you have any health complications during any of your pregnancies, such as hypertension, preeclampsia, or gestational diabetes?

◦ If yes, which complications did you have?
◦ If yes, for how many pregnancies?
6. Did you have any complications during your deliveries, such as postpartum hemorrhaging or the need for a C-section or induced labor?

◦ If yes, for how many pregnancies?
◦ If yes, which complications did you have?
7. For each baby:

◦ Was your baby a boy or a girl?
◦ Was your baby born early or at full term?
◦ What was your ‘due date’ and your baby’s birthday?
◦ Do you know your baby’s weight at birth?
◦ Do you know your baby’s length and head circumference at birth?
◦ Did your baby have any birth defects?
◦ Did your baby need to go to a newborn intensive care unit?

∎ If yes, what was the reason?
◦ Was your baby healthy at the following ages:

∎ 6 months?
∎ 1 year?
∎ 5 years?
∎ [No responses included self-reported reasons, including death of the child]
◦ How old is your child now?

